# Exploring the phytobeneficial and biocontrol capacities of endophytic bacteria isolated from hybrid vanilla pods

**DOI:** 10.1101/2023.02.24.529991

**Authors:** Guillaume Lalanne-Tisné, Bastien Barral, Ahmed Taibi, Zana Kpatolo Coulibaly, Pierre Burguet, Felah Rasoarahona, Loic Quinton, Jean-Christophe Meile, Hasna Boubakri, Hippolyte Kodja

## Abstract

Few studies have been conducted on endophytic bacteria of vanilla. In this study, 58 bacterial strains were isolated from two hybrid vanilla plants from Madagascar, *Manitra ampotony* and *Tsy taitra*. They were genetically characterised and divided into four distinct phylotypes. A selection of twelve strains corresponding to the identified genetic diversity were tested *in vitro* for four phytobeneficial capacities: phosphate solubilisation, free nitrogen fixation, phytohormone and siderophore production. They were also evaluated *in vitro* for their ability to biocontrol the growth of the vanilla pathogenic fungi, *Fusarium oxysporum f. sp. radicis vanillae* and *Cholletotrichum orchidophilum*. Bacteria belonging to three different phyla were found to be highly competent in each of the phytobeneficial capacities tested. Bacteria belonging to the phylum related to *Bacillus siamensis* showed the best capacity to inhibit fungal growth making them good candidates for controlling fungal diseases of vanilla. This competence was highlighted with spectral imaging showing the production of lipopeptides by the bacterial strains confronted with the pathogenic fungi of vanilla.

## 1. Introduction

Vanilla is the only orchid of significant economic importance as an edible crop. It is a tropical hemi-epiphytic plant that grows in a humid environment [1]. About 100 species definethe genus *Vanilla* in the family *Orchidaceaes* [2] but only three species are specifically cultivated for their pods: *Vanilla planifolia* which represents the dominant cultivated species [3], *Vanilla pompona* [4] and *Vanilla tahitensis* [5]. In the 1990s, hybrids were created and disseminated in the cultivation areas of Madagascar: *Manitra ampotony* (*V. planifolia* x *V. tahitensis*) and *Tsy taitra* (*V. planifolia* x *V. pompona*) [6], [7]. *Manitra ampotony* has a high vanillin content and, unlike *V. planifolia*, mature pods do not show a dehisence slit. Thus drying and ripening stages of the pod can take place on the plant and do not require the numerous processing steps necessary to obtain marketable black pods with *V. planifolia* [6], [7]. The hybrid *Tsy taitra* also has a remarkable aroma potential, a high productivity potential, long pods and high resistance to various diseases [8]

Vanilla harbors various endophytic microorganisms which could be used as markers of the terroir [9]. They live asymptomatically in plant tissues for at least part of their life cycle. These endophytic microorganisms (bacteria and fungi) use different mechanisms to exert various beneficial effects from plant growth promotion or biotic and abiotic stresses [10]–[13]. Among endophytic bacteria, a particular class of bacteria called PGPB (Plant Growth Promoting Bacteria) are found in a majority of plants and have specialized and acquired the ability to invade a host [14]. Thus, they are able to fully colonize the rhizosphere and phyllosphere[10], [15], [16]. These endophytic bacteria are involved in physiological mechanisms, such as the production of phytohormones [11], [17] or the stimulation of specific activities of the host plant, leading to an increase in enzyme catalysis, and improving water uptake to facilitate mineral nutrition (ion uptake, siderophore production) and seed germination [18]-[19]. In addition, endophytic microorganisms can occupy an ecological niche that overlaps with that of many plant pathogens, thereby stimulating host plant defenses. Endophytic bacteria are known to inhibit the growth of bacterial and fungal phytopathogens [14] through the production of antibiotics, siderophores and hydrolytic enzymes [20]. These direct benefits may be enhanced by the stimulation of systemic response in the host plant, leading to increased resistance responses against pathogens [16].

Vanilla does not escape fungal attacks. *Colletotrichum orchidophilum*, the agent of black spot disease, is considered as one of the major pathogens of vanilla, attacking the aerial organs of the plant, including the pod, and can be responsible for a reduction in pod production of 10-30% [21]. *Fusarium oxysporum f. sp. radicis-vanillae* is the fungal pathogen responsible for rot and stem rot (RSR), the disease wich has the greatest impact on vanilla production worldwide [22] and has already been isolated from vanilla pods [9]. Bacteria competing in the environment with other micro-organisms possess direct antagonistic activities against pathogens through hyperparasitism or antibiosis and others through competition for nutrients [23], [24]. On the other hand, direct antagonists, which actively produce a broad spectrum of antimicrobial metabolites, are considered most effective against competitors, allowing advantages for antibiotic-producing microorganisms in resource-limited environments [25]. Several studies have been carried out on the phytobenefic properties of orchid endophytes [26]. Nevertheless, few studies have addressed the subject of vanilla endophytes[27]–[29] and none on the bacteria hosted by the vanilla pod, whereas the bacteria isolated from this organ could have remarkable properties. The isolation of bacteria from vanilla pods that possess antagonistic activities against the above-mentioned vanilla pathogens is a crucial step for the development of adequate and effective biocontrol products against these diseases.

The present study aims: i) to isolate and purify endophytic bacteria from green pods of the hybrids *Manitra ampotony* and *Tsy taitra*, ii) to study the diversity of its strains by 16S RNA gene sequencing and phylogenetic analysis, iii) to evaluate their potential effectiveness in promoting plant growth and iiii) to evaluate their antagonistic activities against *Colletotrichum orchidophylum* and *Fusarium oxysporum* f. sp. *radicis-vanillae*.

## 2. Materials and Methods

### 2.1. Isolation of bacteria and condition of growth

Endophytic microorganisms were isolated from mature green pods of two accessions of vanilla hybrids from Madagascar; *Manytra ampotony* and *Tsy taitra*. Three pods for each accession were collected at the same time in june 2014 from three different plants grown close to each other at the vanilla plantation Ambohitsara in Sambava region in north-east Madagascar. The epidermis of the pods were sterilized using the protocol of Khoyratty et al. (2015) [9]. Four 3–5 mm thick slices of sterilized pod were placed in a Lysogenic Broth (LB) medium agar plate (1.5% agar). Colonies were subcultured several times on a fresh LB agar plate until single colonies were obtained. Bacterial strains were stored at −80°C in cryotubes (Mast cryobank, United Kingdom).

### 2.2. DNA extraction

For each strain, a bacterial pellet of 48h-bacterial cultures (10 μL) was resuspended in 100 μL of RES buffer (Macherey Nagel™) and supplemented with lysozyme (20 mg/mL). After shaking and incubation for 10 minutes at room temperature, 200 μL of sodium dodecyl sulfate (0.1%) and 300 μL of phenol-chloroform-isoamyl alcohol (25:24:1) were added. The suspension was homogenized then centrifuged at 18000 g and 5 °C for 5 minutes. The aqueous phase was recovered and then supplemented with 300 μL of phenol-chloroform-isoamyl alcohol (25:24:1). After shaking and centrifugation at 18000 g and 5 °C for 10 min, the aqueous phase was then supplemented with 1/10 of its volume of sodium acetate and 1 volume of isopropanol. The suspension was mixed by turning it several times and stored at −20 °C for 15 hours. This suspension was centrifuged for 30 minutes at 18000 g and 5°C then the supernatant was removed. The DNA pellet was dried for a few minutes in the open air and resolubilized in 100 μL elution buffer (TRIS-HCl pH 8.5). The concentration and quality of the extracted DNA was estimated by Nanodrop^®^ spectrophotometer (Thermo Scientific NanoDrop™ 1000 Spectrophotometer). The bacterial DNA of the 58 strains was normalized to 20 ng.μL^-1^.

### 2.3. PCR amplification and sequencing of 16S rRNA gene

PCR amplification of the 16S rRNA gene was performed using the bacterial universal primers: pA (5′-AGAGTTTGATCCTGGCTCAG-3′)/pH (5′-AAGGAGGTGATCCAGCCGCA-3′) [30] and 27F (5′AGAGTTTGATCMT-GGCTCAG-3′) [31] / 1378R (5′-CGGTGTGTACAAGGCCCGGGAACG-3′) [32].

Additional PCR with primers targeting the internal region of the 16S rRNA gene - COM1 (5′- CAGCAGCCGCGGTAATAC-3′)/COM2(5′-CCGTCAATTCCTTTGAGTTT-3′) [33], [34] was performed in case we obtained poor quality or short sequences. PCR reactions were conducted in a total volume of 25 μl containing 2.5 μL of buffer (10X), 0.75 μL of MgCl_2_ (50 mM), 0.5 μL of dNTPs (10 mM), 2.5 μL forward primer (10 μM), 2.5 μL reverse primer (10 μM), 2.5 μL of Dimethylsulfoxide (DMSO), 0.125 μL of Taq polymerase (Promega), 12.625 μL of H_2_O and 1 μL bacterial DNA (20 ng/ μL). PCR reactions were incubated for 4 min at 94 °C followed by 30 cycles of [50 s at 94 °C, 50 s at 55 °C and 90 s at 72 °C] with a final extension of 3 min at 72 °C. PCR products were visualized by agarose gel electrophoresis (1% in TAE buffer) after Ethidium Bromide Staining. The PCR fragment was then purified on a Nucleobond AX10000 column (Macherey NagelTM) and concentration of DNA recovered were determined with a spectrophotometer (Thermo Scientific NanoDrop™ 1000 Spectrophotometer). The PCR products were sequenced by Microsynth France SAS using same primers.

### 2.4. Sequence analysis and taxonomic identification

DNA chromatograms were checked using SnapGene^®^ Viewer 6.0.2 software and the assembly was performed with CAP3 software [35]. The sequences obtained were aligned with the EzBioCloud Database Update 2021.07.07 [36]. Alignments were performed using ClustalW, and phylogenetic trees were constructed using the “Maximum Likelihood” method associated each time with a bootstrap of 500 repetitions in Molecular Evolutionary Genetics Analysis (MEGAX) software [37]. All sequences were submitted to GeneBank^®^ database.

### 2.5. Phosphate Solubilizing Efficiency

The phosphate solubilizing efficiency of the selected strains was determined by following the protocol of Mehta and Nautiyal [38], [39]. In brief, bacterial strains were grown for 7 days at 28 °C with continuous agitation (140 rpm) in NBRIP-broth medium containing Bromophenol blue (BPB) as pH indicator. *Pseudomonas fluorescens* (F113), a strain known as plant growth-promoting agent [40], [41] was used as a positive control and non-inoculated medium served as a negative control. All these experiments were reproduced at least three times with three replicates. At the end of the incubation period, the final OD_600_ values were subtracted from the negative control values with the following equation :

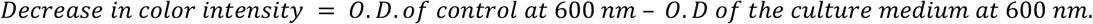

### 2.6. Siderophore production

The selected bacterial strains ability for siderophore-producing was verified by universal Chromo Azurol S (CAS) assay [42]. Prior to the experiment, glassware was cleaned with 3 mol/L hydrochloric acid (HCl) to remove iron and subsequently washed in deionized water [43]. The 24 h-bacteria cultures were used as inoculum, and bacterial concentrations were adjusted so that the optical density at 600 nm (OD600) was 0.5 in NaCl solution (0.85%). Then, 10 μL of each suspension was spotted onto a modified double-layered chromo azurol S (CAS) - MM9 agar plate[44]. MM9 agar as the bottom layer served as nonselective rich medium for bacterial growth. To measure the Fe-chelating function of siderophores, the experiments were performed with CAS-blue agar (dye solution 10 mL) as the top layer. A CAS reagent was prepared according to Schwyn and Neilands[42]. Briefly, 121 mg CAS was dissolved in 200 ml distilled water and 10 ml of 1 mM ferric chloride (FeCl_3_,6 H2O) solution prepared in 10 mM HCl. Then, 145.8 mg of Hex-Decyl Tri-Methyl Ammonium bromide [HDTMA] was slowly added with constant stirring to the resulting dark purple mixture. The pH of the resulting dark blue solution was adjusted to 5, then 6.048 g of Piperazine-N, N’-bis ethane sulfonic acid (PIPES) was added slowly with constant stirring to mixture. While stirring, the pH was adjusted to 6.8. The Chromo Azurol S (CAS) agar (0.9 % w/v) solution was overlaid onto colonies of solid culture. All reagents in the indicator solution were freshly prepared for each batch of CAS-agar. The plates were incubated for 7 days at 28 °C. Sterile water was used as a negative control, and *Pseudomonas fluorescens* F11 was used as a positive control. Experiments were independantly reproduced at least three times each with three replicates. The appearance of an orange halo in the CAS-agar was evaluated (Figure S1). The percentage of halo was determined according to Pinter et al., 2017 [45] by the equation:

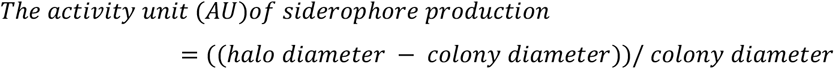

### 2.7. Indole-3-acetic acid production by using Salkowski’s reagent

To evaluate the capacity of bacterial strains to produce indole acetic acid (IAA), the production levels were determined on Yeast Extract Mannitol broth medium (YEM) amended with L-Tryptophane

(2mg.mL^-1^) and KNO_3_ (1%) according to Vincent method [46] with modifications. The broth medium was inoculated in triplicate with 24 h-bacteria cultures, and bacterial concentrations adjusted so that the optical density at 600 nm (OD600) was 0.1. The cultures were incubated in dark for 7 days at 28.0 °C, and then 1 mL of each culture was centrifuged at 5000 rpm for 30 min. Supernatant (100 μl) of each bacterial culture was added in separate wells of microplate followed by the addition of 100 μl of Salkowski’s reagent [50 mL perchloric acid 35% HClO_4_ and 1 mL of 0.5 M ferric chloride FeCl_3_] and incubated at 37 °C for 30 min. A pink coloration confirmed the presence of IAA in the supernatant, which was quantified using spectrophotometer at 535 nm. The cultures in YEM broth without L-Tryptophane and KNO_3_ was used as controls. Sterile water was used as a negative control, and Pseudomonas fluorescens (F113) was used as a positive control.

### 2.8. Bacterial anaerobic growth in nitrogen-free solid medium

To determine the nitrogen (N) fixing capacity of selected strains, the 24 h-bacteria LB cultures were centrifuged and washed with NaCl 0.85 % (3 times). Bacterial concentrations were adjusted to an optical density at 600 nm (OD600) of 0.1 in NaCl solution, then incubated in vials with semi-solid NFb medium [47], [48] (components per liter: 5 g malic acid, 0.5 g K_2_HPO_4_, 0.1 g NaCl, 0.2 g MgSO^4^·7H_2_O, 0.02 g CaCl_2_·2H_2_O, and 2 mL of bromothymol blue (0.5% in 0.2 N KOH solution; pH indicator), 4.5 g KOH (pH control)), supplemented with 2 mL micronutrient solution (components per 100 mL: 0.004 g CuSO_4_·5H_2_O, 0.12 g ZnSO_4_·7H_2_O, 0.14 g H_3_BO_3_, 0.1 g Na_2_MoO_4_·2H_2_O, 0.117 g MnSO_4_·H_2_O), 1 mL of vitamin solution (0.1 g.L^−1^ biotin, 0.2 g.L^−1^ pyridoxine) and 4 mL of Fe-EDTA solution (1.64 % (w/v)) and solidified with 1.8 g Oxoid agar-agar, pH was adjusted to 6.5 – 6.8 with NaOH. Each plate was incubated for 5 to 7 days at 30 °C. The non-inoculated medium served as a control. Nitrogen fixing capacity is visually evaluated by the presence and the thickness of the formation.

### 2.9. Antagonism assay

The antifungal activity of the selected bacterial strains was evaluated *in vitro* against two pathogens of Vanilla, *Colletotrichum orchidophilum* (BS11) [21] and *Fusarium oxysporum sp. Vanillae* (Fo72a) [22]. For the confrontation test, a LB liquid preculture of each bacterial strain was prepared during 24 hours at 33°C under mild shaking. The concentrations of 24h-LB bacterial culture were estimated by calculating the OD at 600 nm, then each preculture was diluted to an OD_600_ of 0.1. The two fungi were grown in PDA medium petri dishes. A 0.5 cm diameter chunk of agar containing a one-week culture of fungus was placed on the center of a PDA agar plate and 5 μL of the diluted bacterial preculture was deposited at 2 cm from the fungus. A negative control with water and a positive control with 5 μL of cycloheximide (100 mg.mL^−1^) were realized. Plates were incubated at 27 °C for seven days. To estimate the inhibition rate of each bacterium against fungus, this formula was used :

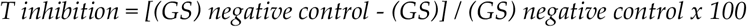

where the growth surface (GS) was estimated as below :

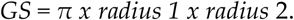

The inhibition zone (IZ) was also measured (figure 1). All distances were estimated in mm with imageJ software [49]. All experiments were indepandatly reproduced three times each with three replicates.

**Figure 1.**
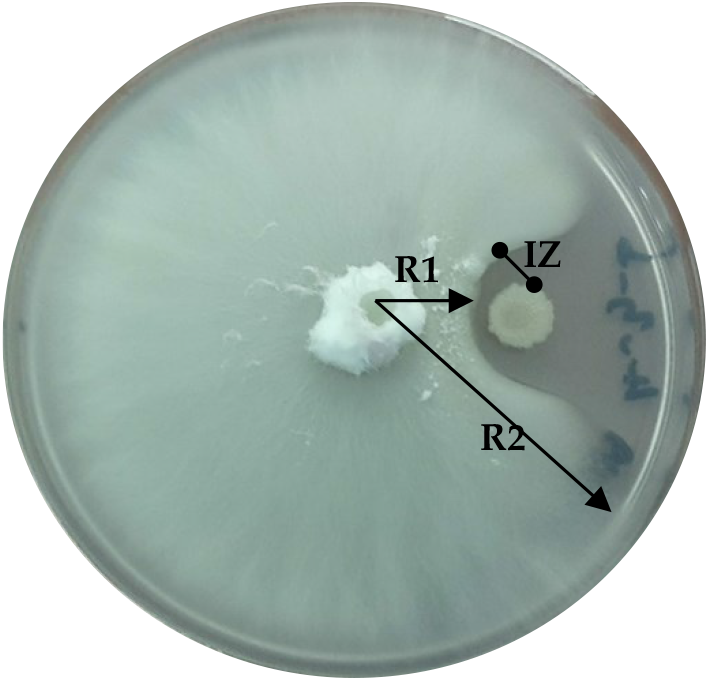
Antagonistic culture of *Colletotrichum orchidophilum* (BS11) with m64 bacterial strain. R1 = radius 1, R2 = radius 2 and IZ = inhibition zone.

### 2.10. Visualization and identification of antifungal metabolites

#### 2.10.1. Mass Spectrometry imaging and data processing

The bacterial strains used for this analyses were m65 and m72 which respectively belong to *Bacillus siamensis* and *Bacillus thuringiensis* species. The fungal strains were *Fusarium oxysporum sp. Vanillae* (Fo72a) and *Colletotrichum orchidophilum* (BS11). Three microliters of both bacterial cell suspension (OD 600nm = 0.1) and fungal suspension (10^5^ conidia.mL^−1^) were spotted at 10 mm distance on a PDA medium diluted to ⅕ enriched with agar to reach a final agar concentration of 7 g.L^−1^. Plates were incubated at 30 °C for 48 h. Microbial samples grown on the agar medium were cut directly from the Petri dish and transferred to a microscope slide (VWR, USA), previously covered with a double-sided conductive copper tape (3M). This assembly was then dried at 39 °C for 2 h. For MSI, a HCCA matrix solution was prepared at 5 mg.mL^−1^ in 80/20 ACN/water doped with 0.1% TFA (Sigma-Aldrich, Belgium). Then two times 10 layers of matrix were sprayed onto the slides with the SunCollect instrument (SunChrom, Germany). The first layer was sprayed at a flow rate of 10 μL.min^−1^. The flow rate was increased by 10 μL.min^−1^ after each layer until it reached 60 μL.min^−1^. MSI samples were acquired on Solarix XR 9.4T (Bruker Daltonics, Bremen, Germany) with a file size of 2M (FWMH ± 300,000 @ 400m/z). The mass spectrometer was systematically mass calibrated from 200 m/z to 2500 m/z before each analysis with a red phosphorus solution in pure acetone spotted directly onto the slides covered with a double-sided conductive copper tape to reach a mass accuracy better than 0.3 ppm. FlexImaging v5 (Bruker Daltonics, Bremen, Germany) software was used for MALDI MS imaging acquisition, with a pixel step size for the surface raster set to 200 μm. For each mass spectrum, one scan of 20 Laser shots was performed at a repetition rate of 200 Hz. The LASER power was set to 14% and the beam focus was set to “small”. Finally, data processing was performed with SCiLS Lab 2016b (SCiLS, Bremen, Germany). Images shown were generated after total ion count normalization.

#### 2.10.2. Liquid Chromatography−Mass Spectrometry Analysis and data processing

Sample for TIMS-TOF analysis were preparate as follows: Agar plugs containing bacteria, confrontation zone and fungus were immersed in 1 mL of acetonitrile/water/trifluoroacetic acid (70/30/0.1 v/v/v). The mix was stirred at 850 rpm and 20 °C for 5 hours (Thermomix.). The supernatant was diluted 1:10 in water before analysis. The chromatographic separation was performed on a M-ClassACQUITY UPLC (Waters, Milford, MA, United States). A 3 min-long sample trapping step was first achieved on a reversed-phase (RP) ACQUITY UPLC M-Class Trap Column (Symmetry C18, 100 Å, 5 μm, 180 μm × 20 mm, Waters, United States) prior to releasing on a ACQUITY UPLC M-Class BEH C18 analytical column (100 Å, 1.8 μm spherical silica, 75 μm × 100 mm, Waters, United States). Water and acetonitrile both supplemented with 0.1% (v:v) of formic acid (FA) were used as eluents and mixed according to a 32 min-long gradient method. The flow rate was set at 0.6 μL.min^−1^. The mass detection was performed on a timsTOF spectrometer (Bruker, Bremen, Germany) with the following parameters: tims Off, scan range m/z, 100–2200. The untargeted profiling data were acquired using Auto MS/MS. Experimental data were processed using Bruker Compass DataAnalysis 6.0.

## 3. Results

### 3.1. Isolation and phylogenetic identification of bacterial strains

The culture of bacterial phenotypes on solid LB medium resulted in 58 different bacterial strains. These strains were associated on the basis of their 16S sequences with species already described with more than 98.5% identity (table 1). The 58 strains are divided into 6 species (*Bacillus subtilis, Bacillus siamensis, Bacillus inaquosorum, Bacillus albus, Bacillus thuringiensis gv. thurigiensis, Curtobacterium oceanosedimentum*). The community of cultivable bacteria isolated from green vanilla pods from *Tsy taitra* (t) consisted of 34 strains associated with three species (*Bacillus subtilis, Bacillus siamensis, Bacillus inaquosorum*). The community of cultivable bacteria isolated from green vanilla pods of *Manitra ampotony* (m) consists of 24 different isolates associated with all 6 species (*Bacillus subtilis, Bacillus siamensis, Bacillus inaquosorum, Bacillus albus, Bacillus thuringiensis gv. thurigiensis, Curtobacterium oceanosedimentum*).

**Table 1:** Identification of the isolated bacterial strains. Strains in bold are those that are investigated in the following study for PGP and biocontrol functions. The prefix ” t ” in the strain name means that it was isolated from a mature green pod of *Tsy taitra* and the prefix “m” from a mature green pod of *Manytra ampotony*

| Strain | GeneBank number | Closest taxonomic relative | % Identity |
| --- | --- | --- | --- |
| **m61** | ON878215 | *Curtobacterium oceanosedimentum* ATCC 31317(T) | 98.68 % (1439 pb) |
| **m62a** | ON878216 | *Bacillus siamensis* KCTC 13613(T) | 99.31 % (1455 pb) |
| **m62b** | ON878217 | *Bacillus siamensis* KCTC 13613(T) | 99.58 % (1443 pb) |
| m63a | ON878218 | *Bacillus siamensis* KCTC 13613(T) | 99.52 % (1453 pb) |
| m63b | ON878219 | *Bacillus siamensis* KCTC 13613(T) | 99..65 % (1443 pb) |
| **m64** | ON878220 | *Bacillus siamensis* KCTC 13613(T) | 99.10 % (1451 pb) |
| **m65** | ON878221 | *Bacillus siamensis* KCTC 13613(T) | 99.59 % (1452 pb) |
| m66 | ON878222 | *Bacillus siamensis* KCTC 13613(T) | 99.52 % (1458 pb) |
| **m67** | ON878223 | *Bacillus inaquosorum* KCTC 13429(T) | 99.72 % (1444 pb) |
| m68 | ON878224 | *Bacillus subtilis* NCIB 3610 | 99.25 % (1459 pb) |
| m69 | ON878225 | *Bacillus subtilis* NCIB 3610 | 99.66 % (1471 pb) |
| m70 | ON878226 | *Bacillus albus* N35-10-2(T) | 99.39 % (1474 pb) |
| m71 | ON878227 | *Bacillus subtilis* NCIB 3610 | 99.04 % (1454 pb) |
| **m72** | ON878228 | *Bacillus thuringiensis* gv. *Thuringiensis* ATCC 10792(T) | 98.84 % (1460 pb) |
| m73 | ON878229 | *Bacillus siamensis* KCTC 13613(T) | 99.1 % (1450 pb) |
| m74 | ON878230 | *Bacillus subtilis* NCIB 3610 | 99.18 % (1458 pb) |
| m75 | ON878231 | *Bacillus subtilis* NCIB 3610 | 98.82 % (1443 pb) |
| m76 | ON878232 | *Bacillus siamensis* KCTC 13613(T) | 98.76 % (1452 pb) |
| m78 | ON878233 | *Bacillus siamensis* KCTC 13613(T) | 99.52 % (1449 pb) |
| m79 | ON878234 | *Bacillus siamensis* KCTC 13613(T) | 99.10 % (1452 pb) |
| m80 | ON878235 | *Bacillus siamensis* KCTC 13613(T) | 98.69 % (1451 pb) |
| m81 | ON878236 | *Bacillus siamensis* KCTC 13613(T) | 99.45 % (1456 pb) |
| m82 | ON878237 | *Bacillus subtilis* NCIB 3610 | 99.38 % (1459 pb) |
| m83 | ON878238 | *Bacillus subtilis* NCIB 3610 | 99.25 % (1457 pb) |
| t1 | ON878239 | *Bacillus siamensis* KCTC 13613(T) | 99.52 % (1450 pb) |
| **t2** | ON878255 | *Bacillus subtilis* NCIB 3610 | 99.31 % (1458 pb) |
| t3 | ON878264 | *Bacillus siamensis* KCTC 13613(T) | 99.31 % (1453 pb) |
| t4 | ON878267 | *Bacillus subtilis* NCIB 3610 | 99.31 % (1451 pb) |
| **t5** | ON878268 | *Bacillus subtilis* NCIB 3610 | 99.31 % (1451 pb) |
| t6 | ON878269 | *Bacillus subtilis* NCIB 3610 | 98.67 % (1460 pb) |
| t7 | ON878270 | *Bacillus siamensis* KCTC 13613(T) | 99.04 % (1459 pb) |
| t8 | ON878271 | *Bacillus subtilis* NCIB 3610 | 99.25 % (1457 pb) |
| t9 | ON878272 | *Bacillus subtilis* NCIB 3610 | 98.83 % (1454 pb) |
| t10 | ON878240 | *Bacillus inaquosorum* KCTC 13429(T) | 99.86% (14443pb) |
| t11 | ON878241 | *Bacillus inaquosorum* KCTC 13429(T) | 99.31 % (1448 pb) |
| t12 | ON878242 | *Bacillus subtilis* NCIB 3610 | 99.04 % (1452 pb) |
| t13 | ON878243 | *Bacillus inaquosorum* KCTC 13429(T) | 99.17 % (1449 pb) |
| t14a | ON878244 | *Bacillus siamensis* KCTC 13613(T) | 99.25 % (1459 pb) |
| t14b | ON878245 | *Bacillus siamensis* KCTC 13613(T) | 99.66 % (1453 pb) |
| t15a | ON878246 | *Bacillus siamensis* KCTC 13613(T) | 99.38 % (1459 pb) |
| t15b | ON878247 | *Bacillus siamensis* KCTC 13613(T) | 99.04 % (1459 pb) |
| t16 | ON878248 | *Bacillus inaquosorum* KCTC 13429(T) | 99.65 % (1440 pb) |
| t16b | ON878249 | *Bacillus inaquosorum* KCTC 13429(T) | 99.17 % (1452 pb) |
| t17a | ON878250 | *Bacillus inaquosorum* KCTC 13429(T) | 99.45 % (1450 pb) |
| t17b | ON878251 | *Bacillus subtilis* NCIB 3610 | 99.25 % (1459 pb) |
| t18 | ON878252 | *Bacillus siamensis* KCTC 13613(T) | 99.31 % (1458 pb) |
| t19a | ON878253 | *Bacillus subtilis* NCIB 3610 | 99.8.9 % (1450 pb) |
| t19b | ON878254 | *Bacillus siamensis* KCTC 13613(T) | 98.84 % (1460 pb) |
| t20 | ON878256 | *Bacillus siamensis* KCTC 13613(T) | 99.52 % (1460 pb) |
| t21 | ON878257 | *Bacillus siamensis* KCTC 13613(T) | 99.38 % (1459 pb) |
| t22 | ON878258 | *Bacillus siamensis* KCTC 13613(T) | 99.38 % (1457 pb) |
| t23 | ON878259 | *Bacillus subtilis* NCIB 3610 | 99.41 % (1353 pb) |
| t24 | ON878260 | *Bacillus siamensis* KCTC 13613(T) | 99.25 % (1457 pb) |
| t27a | ON878261 | *Bacillus siamensis* KCTC 13613(T) | 99.52 % (1460 pb) |
| t27b | ON878262 | *Bacillus siamensis* KCTC 13613(T) | 99.31 % (1458 pb) |
| t29 | ON878263 | *Bacillus siamensis* KCTC 13613(T) | 99.38 % (1459 pb) |
| t30 | ON878265 | *Bacillus inaquosorum* KCTC 13429(T) | 99.18 % (1459 pb) |
| t31 | ON878266 | *Bacillus siamensis* KCTC 13613(T) | 99.45 % (1455 pb) |

The 16S rRNA gene sequences were used to reconstruct the phylogeny of the 58 isolates. The resulting phylogenetic tree shows that the populations are divided into 4 phylotypes (Figure 2). Phylotype 1 is associated with the species *Curtobacterium oceanosedimentum*. Phylotype 2 is associated with the species *Bacillus thuringiensis gv. thuringiensis* and *Bacillus albus*. Phylotype 3 is associated with the species *Bacillus subtilis* and *Bacillus inaquosorum* and finally, phylotype 4 is associated with the species *Bacillus siamensis*. The species composing the community of cultivable bacteria isolated from *Manitra ampotony* green vanilla pods belong to all four phylotypes whereas those composing the community of bacteria isolated from *Tsy taitry* mature green pods belong only to phylotypes 3 and 4.

**Figure 2.**
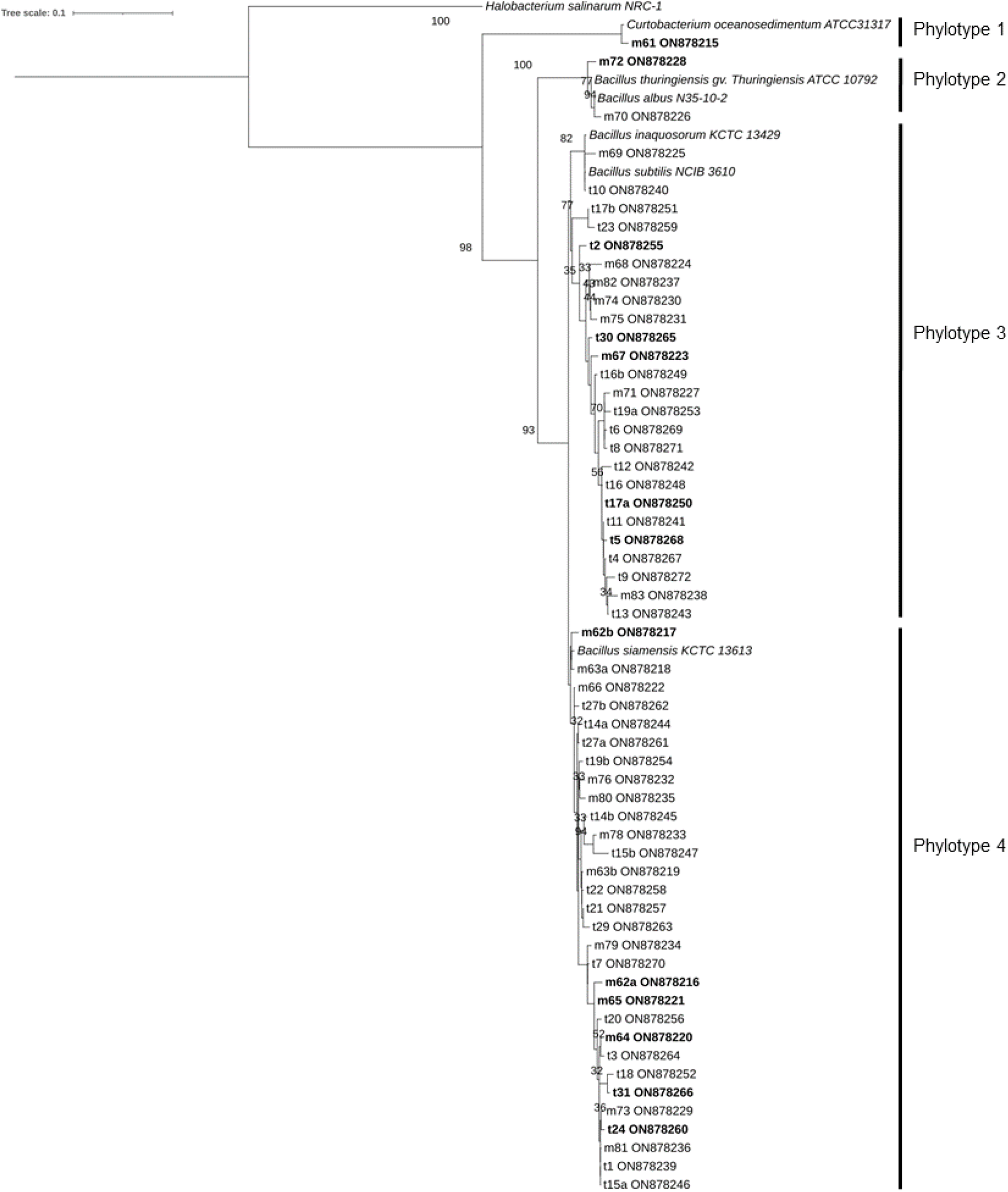
16S rDNA-based dendrogram showing the phylogenetic relationships between the different isolates and the most phylogenetically related species. The tree was rooted with the 16S rDNA sequence of *Halobacterium salinarum* as a reference outgroup. Only ML bootstraop branches that support values greater than 30% are shown

### 3.2. Screening of PGPB capacities

12 strains were selected for further study corresponding to the diversity characterised above. All 12 bacterial isolates excepet m61 did belong to the genus *Bacillus*. M61 isolate is identified as belonging to the genus *Curtobacterium* (table 1) and is the only representative of phylotype 1 (figure 2). The other 11 are distributed in the three other phylotypes as follows: m72 associated with *B. thurigensis* represents phylotype 2; m62a, m62b, m64, m65 and t24 associated with *B. siamensis* represent phylotype 4 and m67, t2, t5, t17a and t30 which are associated with *B. subtilis* and *B. inaquosorm* in phylotype 3. Regarding the capacity to solubilize phospate, the three phylotype 4 isolates (*B. siamensis)* m62a, m64 and m65 showed the highest phosphate solubilisation index value based on Tuckey’s HSD Test for multiple comparisons at p < 0.05. The phylotype 1 isolate m61, had a negligible solubilisation index. For the capacity to produce siderophore, m61 did not produce siderophore. m72, representing phylotype 2 (*B*.*thurigensis)*, had a lower siderophore production than the control *Pseudomonas fluorescens* F113 (0.40 ± 0.27 and 1.29 ± 0.07 AU, respectively). The other isolates studied belonging to phylotype 3 and 4 (*B. siamensis* and *B. subtilis/inaquosorum*) showed a slightly lower level of siderophore production when compared to the control except for t2 which showed a similar value based on Tuckey’s HSD Test for multiple comparisons at p < 0.05. For the production of IAA all studied isolates produced IAA at a similar rate exept m72, representing phylotype 2 associated with *B. thurigensis*, which had the highest rate production of IAA of the study with a OD_535_ of 1.27 ± 0.25.

Finally, concerning the capacity of the strains to grow on a nitrogen-free medium and thus having the capacity to fix atmospheric nitrogen, only m62b isolate did not grow. t17a and t5 isolates associated with phylotype 3 showed the highest capacity to grow.

### 1.1. Antagonism assay

The twelve strains were challenged in antagonistic cultures during seven days with two fungal pathogens of vanilla, *Colletotrichum orchidophilum* (BS11) and *Fusarium oxysporum sp. Vanillae* (Fo72a). Figure 3A shows the growth inhibition rates of *Colletotrichum orchidophilum* (BS11) induced by each bacterial strain co-cultured with the fungus after seven days. A one-way ANOVA was performed to compare the effect of the 12 strains on the growth of BS11 and revealed a statistically significant difference in inhibition rate of the different isolates (p < 2e-16). Tukey’s HSD Test for multiple comparisons revealed that the mean value of inhibition rate score was significantly different between two groups at p < 0.05. In this way, m61, m72, m67, t17a and t30 strains are grouped together as the negative control as they did not show any significant inhibition of the growth of strain BS11; the rate of inhibition in this group is, apart from outliers, between −8.21% and 16.97%. Strains t2, t5, m62a, m62b, m65, m64, and t24 are grouped as the positive control and therefore induced a clear inhibition of the growth of the fungus; the inhibition rate in this group, excluding outliers, was between 56.32% and 83.33%. All strains belonging to phylotype 4 (genus *Bacillus*) showed the ability to inhibit the growth of BS11. The strains m61 and m72 belonging to phylotype 1 and 2 respectively did not shown any ability to inhibit the growth of BS11. However, the strains belonging to phylotype 3 are distributed in the two (negative and positive) groups; m67, t17a and t30 did not show any capacity to inhibit the growth of BS11. On the contrary, t2 and t5 with a rate of inhibition of the growth of the fungus ranging between 56.32% and 67.64%.

**Figure 3.**
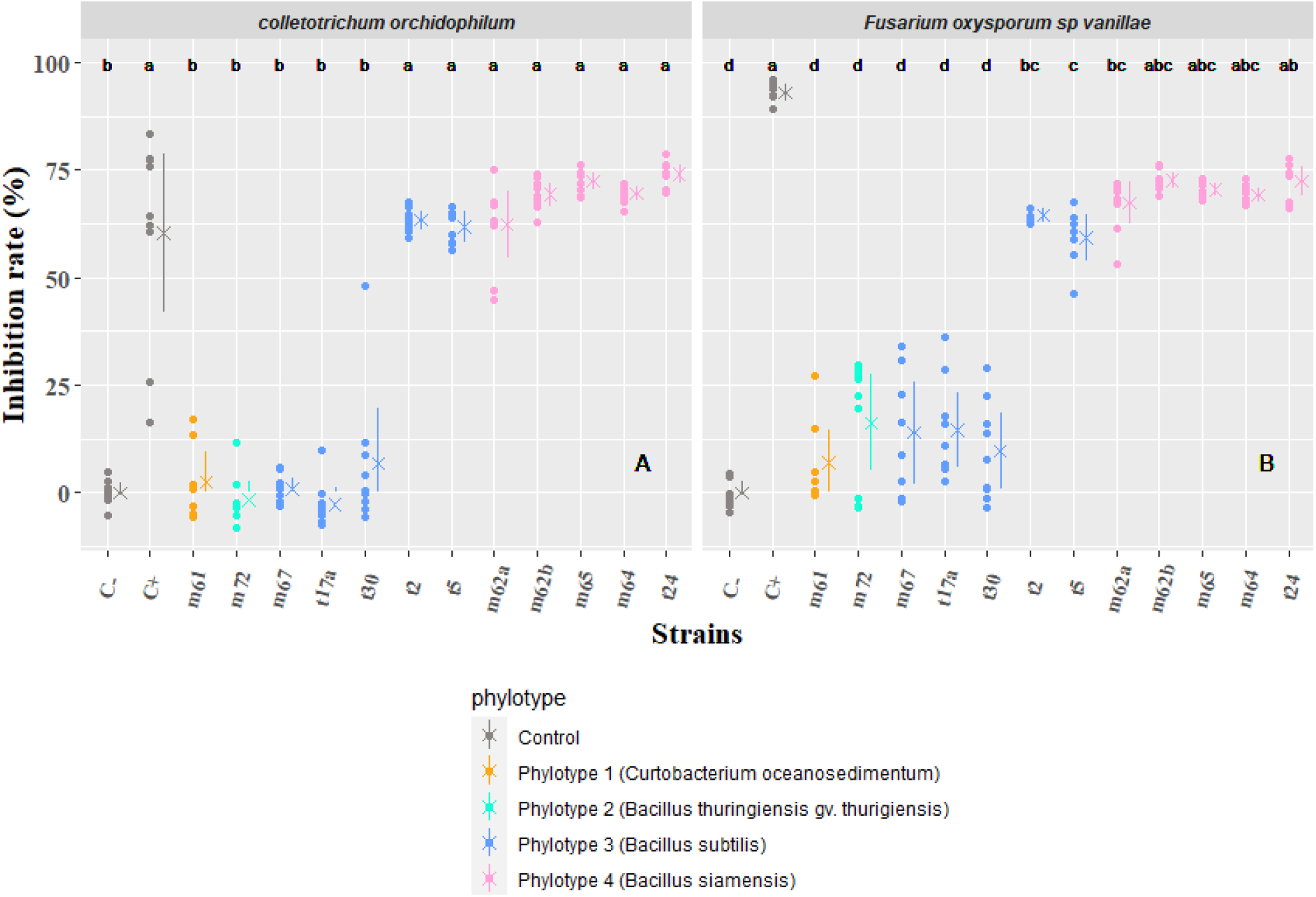
*In vitro* inhibition rate induced after 7 days by each bacterial strain on PDA medium on fungal growth of: (A) *Colletotrichum orchidophilum* (BS11) and (B) *Fusarium oxysporum sp. Vanillae* (Fo72a). Each phylotype is defined by a colour code. Positive (C−: growth of the fungus without confrontation) and negative (C+: cycloheximide) controls appear in grey. A dot is assigned to each measurment. Mean values are represented by crosses and segments indicates standard errors. Different letters indicate that data are significantly different at p < 0.05 between strains (according to Tukey’s multiple comparison test).

The confrontations of the 12 bacterial strains against *Fusarium oxysporum sp. Vanillae* (Fo72a) during seven days (figure 3B) showed a similar pattern as described above for *C. orchidolphilum* (BS11). A one-way ANOVA was performed to compare the effect of the 12 strains on the growth Fo72a and revealed that there was a statistically significant difference in inhibition rate of the different strains (p < 2e-16). Tukey’s HSD Test for multiple comparisons found that the mean value of inhibition rate score was significantly different between more than two groups at p < 0.05. As for the confrontation against *C*.*orchidophilum* (BS11), strains m61, m72, m67, t17a and t30 are grouped in the negative control as no significant inhibition of the growth of strain Fo72a was observed. The positive control showed an extremely high inhibition rate (between 89.04% and 96.24%) ; *Fusarium oxysporum sp. Vanillae* (Fo72a) was more sensitive to the antifungal action of cycloheximide than *Colletotrichum orchidophilum* (BS11). There was a continuum of groups ab, abc, bc and c grouping strains with significant ability to inhibit the growth of Fo72a namely t2, t5, m62a, m62b, m65, m64 and t24. Interestingly, the same strains were identified as able to inhibit both fungal pathogen growth with a similar rate apart from t2, and t5 (phylotype 3) and m62a (phylotype 4) strains which showed a slightly lower inhibition rate (groups bc and c) on *F. oxysporum sp. Vanillae* when compared to *C. orchidophilum*.

The inhibition zone between the bacterial and fungal colonies was measured for bacterial strains that were found to be competent to inhibit the growth of the fungus (figure 4). Figure 4A shows the inhibition zones between the bacterial strains and *Colletotrichum orchidophilum* (BS11) on PDA after seven days of co-culture. A one-way ANOVA was performed to compare the mean of inhibition zone between the bacterial strains and BS11 and revealed that there was a statistically significant difference in inhibition rate of the different strains (p < 2e-16). Tukey’s HSD Test for multiple comparisons found that the mean value of inhibition rate score was significantly different between two groups at p < 0.05. The strains m62b, m64 and t24 are included in the same group and the inhibition zone in this group is, apart from outliers, between 1.11 and 3.03mm. The other bacterial strains were grouped together in the same group which did not induce a zone of inhibition against BS11. The strains m62b, m64, t24, m62b and m65 belonged to phylotype 4 associated with *Bacillus siamensis* but only m62b, m64 and t24 produced an inhibition zone in co-culture with BS11. Even if t2 and t5 strains (phylotype 3) associated with *B. subtilis* and *B. inaquosorum* could inhibit the growth of BS11 they did not produce any inhibition zone. The same trend is observed when bacterial strains are confronted with *Fusarium oxysporum sp. Vanillae* (Fo72a) (figure 4B). A one-way ANOVA was performed to compare the mean of inhibition zones and revealed that there was a statistically significant difference in inhibition rates of the different strains (p < 2e-16). Tukey’s HSD Test for multiple comparisons showed that the mean value of inhibition rate score was significantly different between more than two groups at p < 0.05.

**Figure 4.**
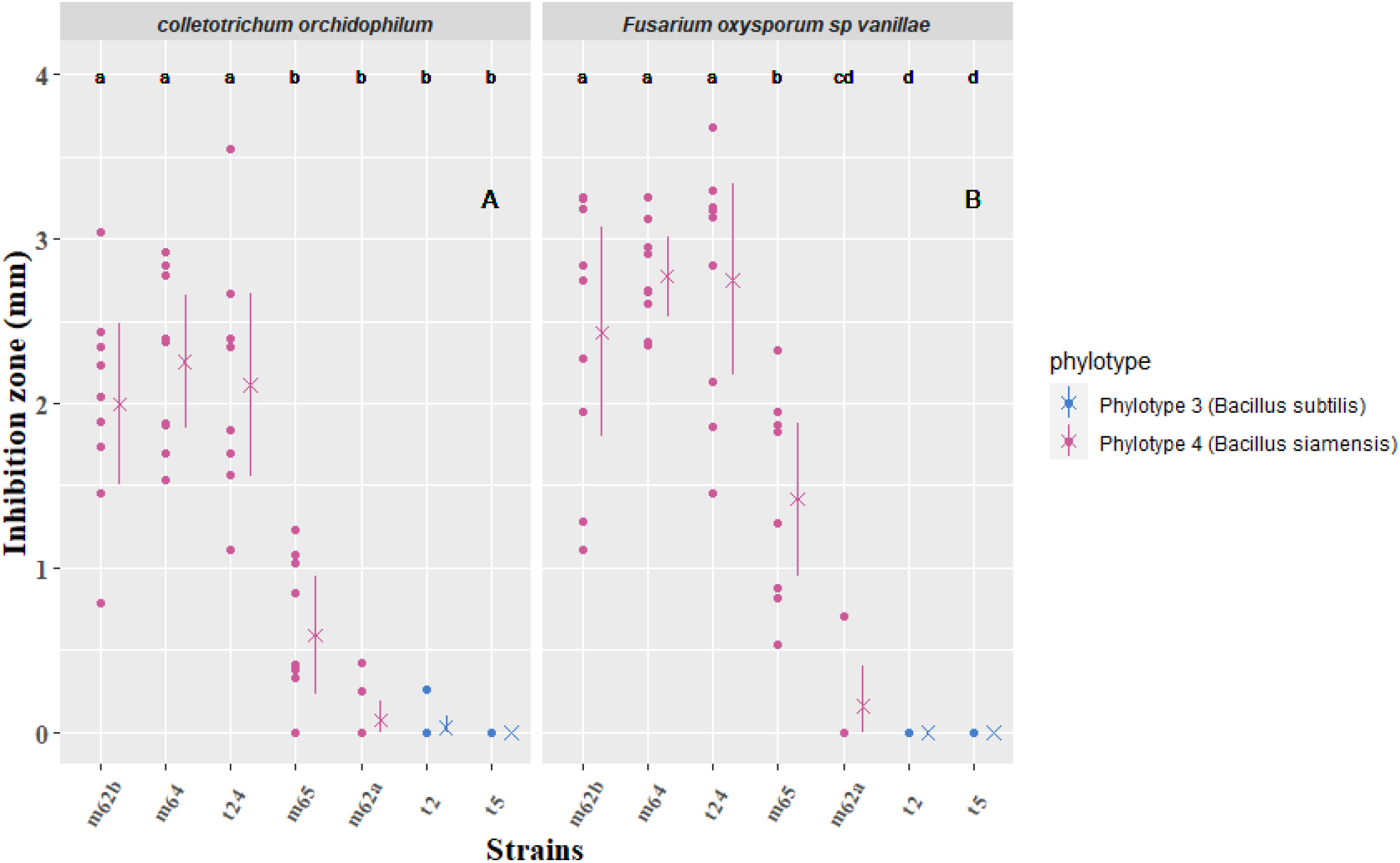
Mean inhibition zone size (mm) measured after seven days of co-cultures between the bacteria and: (A) *Colletotrichum orchidophilum* (BS11) and (B) *Fusarium oxysporum sp. Vanillae* (Fo72a). Each bacterial phylotype appear with a colour code. In grey are the positive (C−: growth of the fungus without confrontation) and negative (C+: cycloheximide) controls. A dot is assigned to each measurment. Mean values are represented by crosses and segments indicate standard errors. Different letters indicate that data are significantly different at p < 0.05 between strains (according to Tukey’s multiple comparison test).

As with BS11, m62b, m64 and t24 strains co-cultured with Fo72a showed the largest inhibition zones with a minimum of1.1mm and a maximum of 3.68mm. t2 and t5 were the only strains (phylotype 3) showed no inhibition zone. Strain m65, with an average of 1.42 ±0.6 mm, showed a more marked inhibition zone against Fo72a when compared to BS11. Strain m62a shows, apart from the outlier, no zone of inhibition in co-culture with Fo72a.

### 1.1. Identification and distribution of bioactive lipopeptides

The chromatographic analysis of *B. siamensis* strain (m65) revealed the presence of three families of lipopeptides (Figure 5A). Mass spectra of ions from each family displayed similar peptide sequences but with varying aliphatic chain lengths. The precursor ions with m/z masses of 994.6440, 1008.6596, 1022.6743 and 1036.6912 corresponded to C12-surfactin, C13-surfactin, C14-surfactin and C15-surfactin, respectively. The fragmentation of the precursor ions allowed for the identification of the amino acid sequence of the surfactins, as confirmed by the peptide sequence Leu-Leu-Val-Asp-Leu-Leu (Figure 5D). Five compounds belonging to the iturin family were identified: iturin A–1 (m/z 1029.5372), iturin A-2 (m/z 1043.5532), iturin A–3 (m/z 1057.5691), iturin A–6 (m/z 1071.5841) and iturin A–8 (m/z 1085.6005) (Figure S1). The mass spectrum decomposition at the m/z 1043.5532 ion revealed the two parts of the iturin-specific peptide sequence: Glu-Asp-Tyr-Asn and Glu-Pro-Asn-Ser. The precursor ions with m/z masses of 1449.7905, 1463.8046, 1477.8203 corresponded to C15-fengycin A, C16-fengycin A, and C17-fengycin A, respectively. In contrast, no lipopeptides were detected in the chromatogram of *Bacillus thuringiensis* strain (m72) when grown in monoculture or in co-culture with a fungus. Mass spectrometry imaging made it possible to visualize the spatial distribution of the lipopeptides produced by bacterial strains confronted with the pathogenic fungi of vanilla (Figure 6). The images clearly show a dividing line between *B*.*siamensis* (m65) and the fungi unlike the images of *B. thuringiensis* (m72) where the fungi cover the bacterial strain. The analysis of the mass spectrometry images reported similarities in the spatial distribution of the lipopeptides belonging to the same family which could be classified according to their relative intensity. Two lipopeptides of each family with the highest intensity have been represented (Figure 6A and 6B). The distribution of iturins was diffuse all around the bacterial colony and over the entire fungal growth inhibition zone. The distribution of the fengycin family, produced by *B. siamensis*, was detected in a short halo around the colony of *B. siamensis*. The localization of surfactins varied depending on the fungus against which the bacterium grew. In the case of the confrontation between *B. siamensis* (m65) and *F. oxysporum f. sp. Vanillae* (Fo72a) surfactins distribute around the colony with higher intensity towards the outer parts. Another pattern is obsverved in the confrontation between *B. siamensis* and *C. orchidophilum* (BS11) where the surfactins are polarized on the confrontation zone.

**Figure 5.**
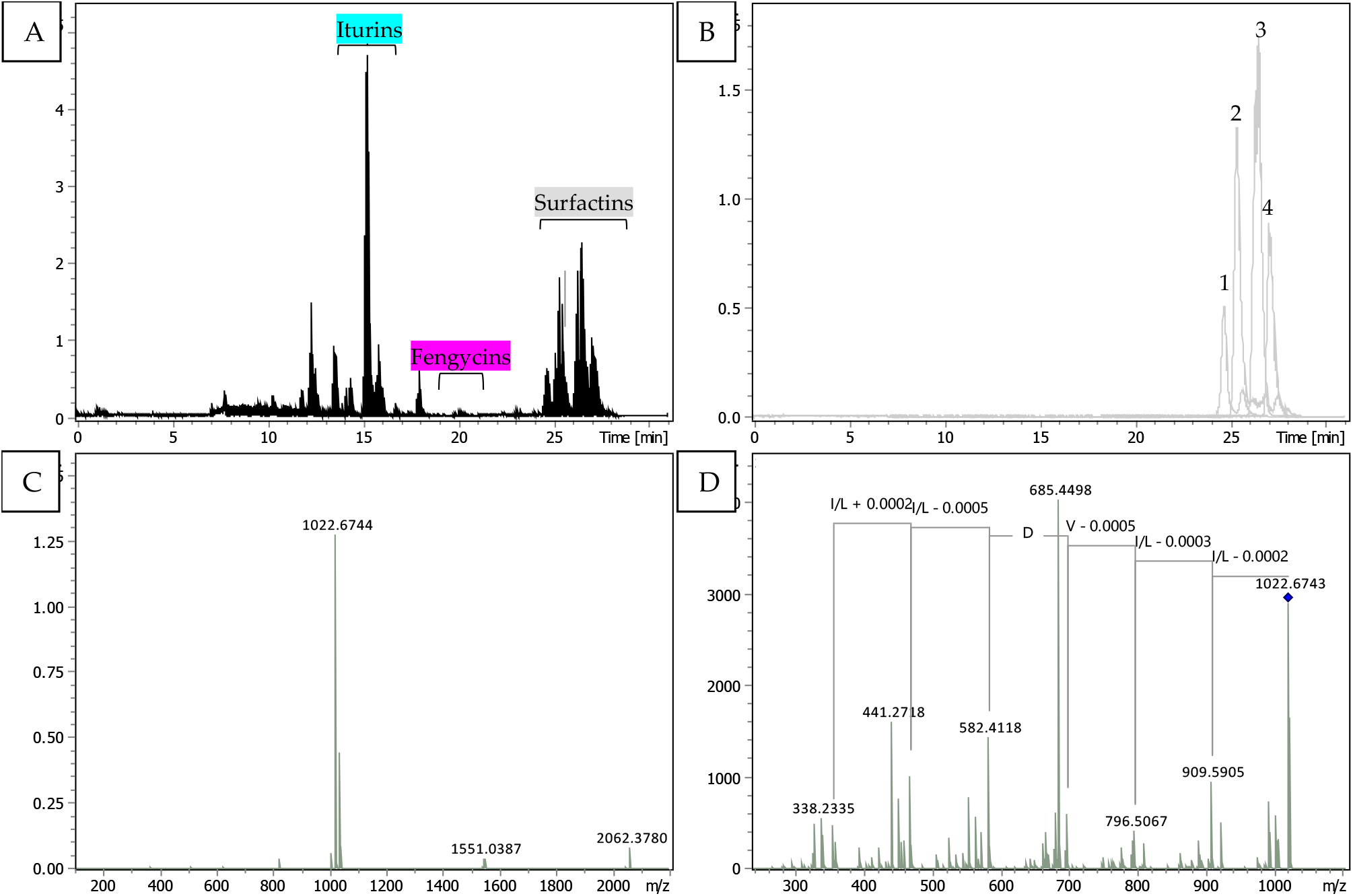
TIMS-TOF analysis of compounds recorded in the inhibition area between *Bacillus siamensis* (m65) and *Fusarium oxysporum sp. Vanillae* (Fo72a). (A) Typical chromatogram with distinct separation times of the three categories of identified lipopeptides. (B) The extracted ion chromatogram of 994.6440 (1), 1008.6596 (2), 1022.6743 (3) and 1036.6912 (4) m/z representing the four present metabolites of surfactins. (C) MS spectrum represented the single charged M+H+ ion of surfactin 1022.67 m/z. (D) MSMS spectrum ion at 1022.67 m/z and the fragmentation lead to the peptide sequence analysis of surfactin. Similar processes of identification for iturins and fengycins families are presented in **Figure S2**.

**Figure 6.**
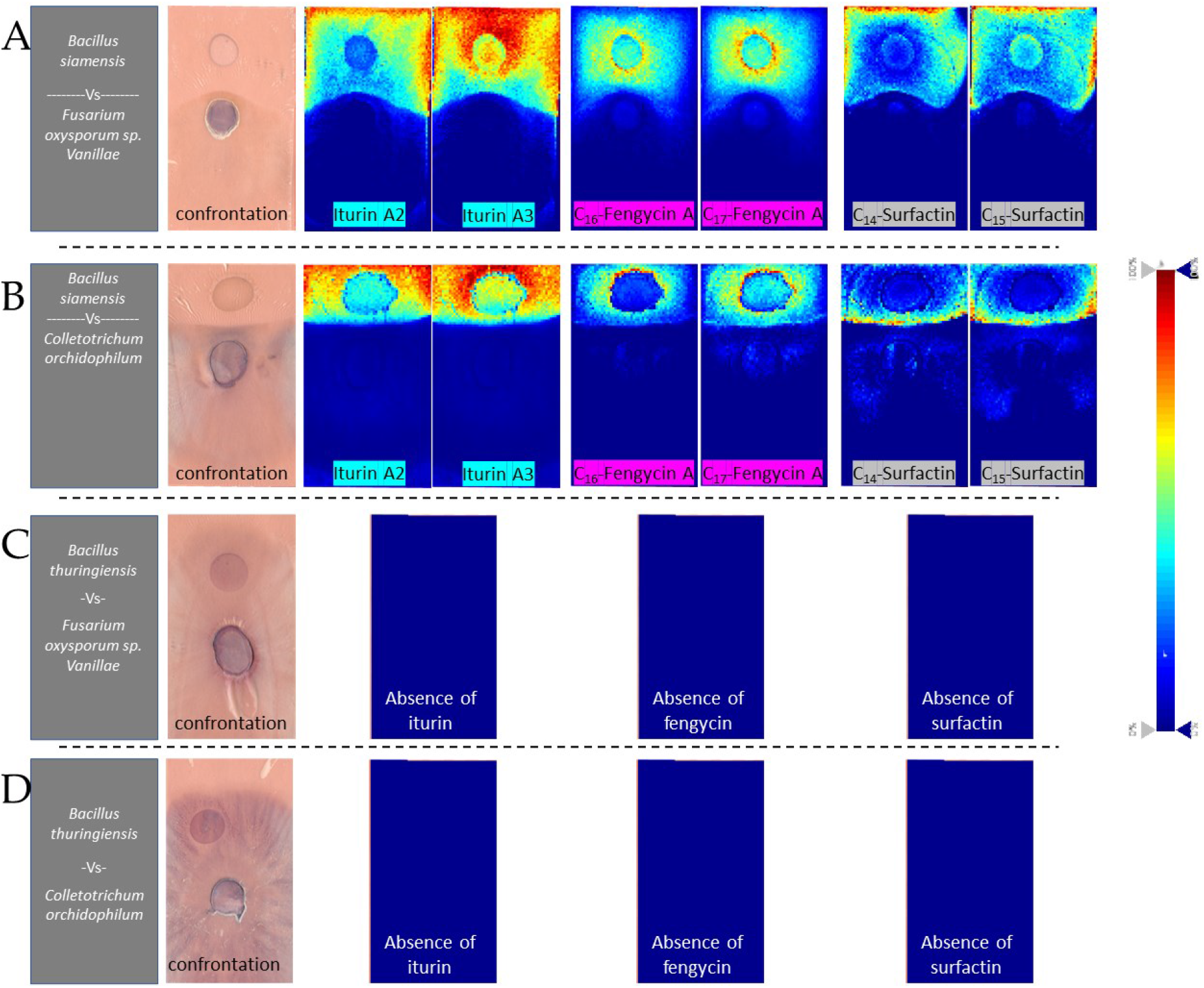
Mass spectrometry images showing the involvement of lipopeptides in *Bacillus spp*. antagonism against pathogenic fungi of vanilla. The images, from left to right, show the confrontation plate with the microorganisms, the distribution of various lipopeptides, and intensity represented by a color gradient. A and B depict *B. siamensis* (m65) against (A) *F. oxysporum sp. Vanillae* (Fo72a) and (B) *C. orchidophilum* (BS11). From left to right; Picture of the confrontation plate, containing the microorganisms in culture transferred to the glass slide, previously covered with double sided conductive copper tape. Distribution of sodium adducts of Iturin A2 (m/z 1065.549, ± 25 mDa), Iturin A3 (m/z 1079.564, ± 25 mDa), C_16_-Fengycin A (m/z 1485.821, ± 25 mDa), C_17_-Fengycin A (m/z 1499.830, ± 25 mDa), C_14_-Surfactin (1044.676, ± 25 mDa) and C_15_-Surfactin (1058.689, ± 25 mDa). (C) and (D) panels show *B. thuringiensis* (m72) against (C) *F. oxysporum sp. Vanillae* (Fo72a) and (D) *C. orchidophilum* (BS11). From left to right; Picture of the confrontation plate, containing the microorganisms in culture transferred to the glass slide, previously covered with double sided conductive copper tape; No lipopeptides were detected in these images. The intensity of each compound is represented by a color gradient ranging from blue to red (0% to 100% intensity).

Mass spectrometry images confirmed the absence of lipopeptides for the strain *Bacillus thuringiensis* (m72) (Figure 6C and 6D).

## 2. Discussion

In this study, we identified a population of endophytic bacteria isolated from the mature pods of two vanilla hybrids and then measured their phytobeneficial potential. The two Vanilla hybrids were *Manitra ampotony* (*V. planifolia x V. tahitensis*) which accumulating high levels of vanillin and *Tsy taitra* (*V. planifolia x V. pompona*) resistant to fungal diseases. Although a greater number of bacterial strains were isolated from *Tsy taitra* pods (34 for a total of 58), they were distributed in only two of the four phylotypes identified, whereas the 24 strains isolated from *Manitra ampotony* pods were distributed in all four phylotypes (table 1). Bacteria strains associated with species of the genus *Curtobacterium* and *Bacillus* of phylotype 2 (*B. thuringiensis* and *B. albus*) were only isolated from pods belonging to *Manitra ampotony*. However, the pods were collected at the same time from hybrids plants grown close to each other at the vanilla plantation Ambohitsara. This suggest a relationship between the composition of the bacterial communities and the genotype of the host plant. It has been shown that there is a close relationship between the microbiome and the genotype of the host plant. This observation is in agreement with different studies [50]–[53] showing that the composition of the bacterial communities is more dependent on the genotype of the host plant than on environmental parameters such as terroir. This indicates that the plant selects its own microbiome.

m61 was the only identified strain of phylotype 1 associated with the *Curtobacterium* genus. Many bacterial species the Actinomycetes class are known to be endophytic bacteria with many PGP traits [54]. However, bacteria of the genus *Curobacterium* are also known to be plant pathogenic bacteria. The best known example is *Curtobacterium flaccumfaciens* which causes disease in many species of *Fabaceae*

[55]–[57]. So far, no studies reported *Curtobacterium* species as pathogen of vanilla or other orchid species. In our study, m61 did not show any particular PGP trait *in vitro* or biocontrol capacity of pathogenic vanilla fungi. It is possible that m61 has a neutralist interaction with the plant host. It is also possible that the LB culture medium used in the study did not favour any optimal development of m61. Indeed, many studies use more specific media for actinomycetes like the ‘Actinomycete Isolation Agar’ (AIA) [58].

In this study, the Phylotype 2 was characterised for its phytobeneficial potential with the m72 strain. The closest taxonomic relative was *Bacillus thuringiensis gv. Thuringiensis. Bacillus thuringiensis*, abbreviated in its commercial form to *Bt*, is well known for its insecticidal properties and is still one of the most widely used biopesticides [61], [62]. It appeared interesting to test a strain close to *B. thurigensis*, for its antifungal capacities and PGP functions. Thus, the m72 strain did not show any antifungal potential against the fungi responsible for the main vanilla diseases. These observations are consistent with the metabolic analyzes where no lipopeptide could be identified in the conditions used in the present study. Regarding its PGP functions, the m72 strain did not show any particular competence in the assimilation of phosphate and nitrogen or in the production of siderophores, but was the best produceder of IAA *in vitro* from tryptophan. It has been shown that IAA-producing rhizobacteria participate favourably in seed germination rate and plant growth [63], [64]. Strain m72 appears to be an interesting candidate for further *in vivo* studies.

Strains of phylotype 3 were associated with *B. inaquosorum* or *B. subtilis* based on 16S sequences similarity (table 1). However, the phylogenetic tree in Figure 2 shows that while the 16S sequences of the reference strains of *B. inaquosorum* and *B. subtilis* were indeed very similar, the strains associated with these two bacteria (phylotype 3) apeared to be relatively distant. Until recently *B. inaquosorum* was considered a subspecies of *B. subtilis*, but consistent genomic and biochemical data have ruled in favour of two different species [59]. This could indicate that the phylotype 3 bacteria could be closely related but distinct species of *B. inaquosorum* and *B. subtilis*. This hypothesis is reinforced by the variability of results observed within the phylotype 3 strains that were selected to test the PGPB traits (table 2). Indeed, the t2 strain showed the highest level of siderophore production with the F113 control, while the t17a and t30 strains of the same phylotype exhibited reduced levels. Similarly, t5 and t17a were the best nitrogen-free fixing strains of the study, while the other strains of phylotype 3 showed a lower level of nitrogen fixation.

**Table 2:**
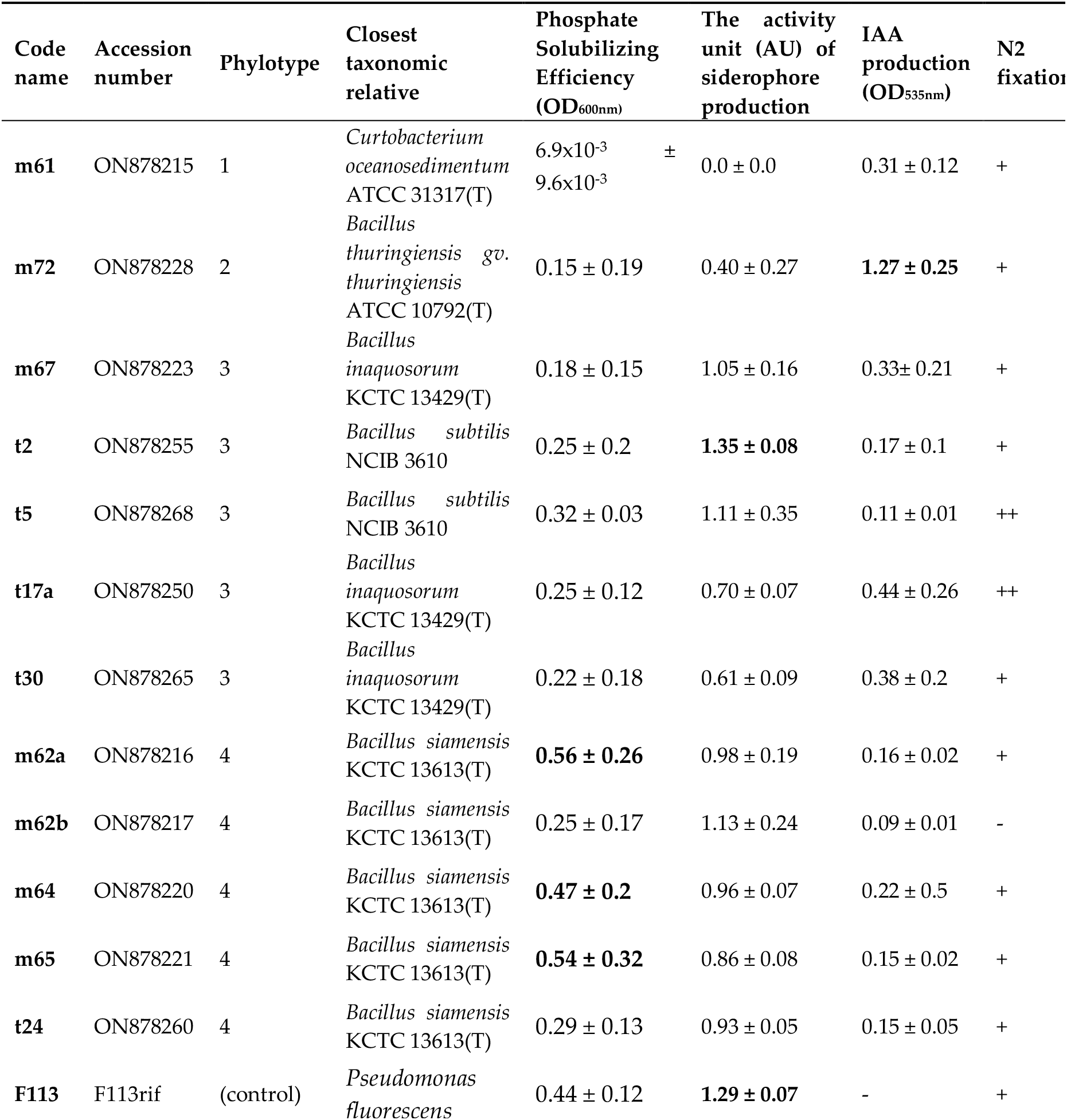
Determination of phosphorus solubilization, IAA production ability, siderophore production and nitrogen fixation capacity of the different strains. Highest values appear in bold highest based on Tuckey’s HSD Test for multiple comparisons at p < 0.05. (−) indicates negative result, (+) and (++) represent nitrogen fixation capacity demonstrated by the presence and thickness of the pellicle formed in nitrogen-free semi-solid media (NFb).

Strains t2 and t5 were the only bacterial strains of phylotype 3 that showed an ability to significantly decrease the growth of *F. oxysporum f. sp. Vanillae* (Fo72a) and *C. oxysporum* (BS11) (figure 3). However, no inhibition zone between bacteria and fungi was observed (figure 4). Although some strains of *B. subtilis* have been shown to be competent as biocontrol agents in other studies [60], [61], strains of phylotype 3 close to *B. subtilis* did not show as much biocontrol capacity in our study.

Bacterial strains belonging to phylotype 4 associated with *Bacillus siamensis* appeared to form a homogeneous group based on both phylogenetic and functional characteristics. These strains produce weak indole acetic acid, fix moderate levels of free nitrogen (with the exception of strain m62b), and produce siderophores. Specifically, m62a, m64 and m65 strains exhibit the highest levels of phosphate solubilization efficiency in comparison to the control strain F113. Additionally, they displayed a higher capacity to inhibit the growth of *Fusarium oxysporum f. sp. Vanillae* (Fo72a) and *C. orchidophilum* (BS11) as evidenced by the clear zones of inhibition observed in confrontation cultures with strains t24, m62b, m64 and m65. To further understand this inhibitory properties, metabolic studies were conducted. The results indicated that the lipopeptides (surfactin, fengycin, and iturin), produced by strain m65 (*Bacillus siamensis*), played a crucial role in the biocontrol of pathogenic fungi [62]. These lipopeptides have been shown to possess broad-spectrum antifungal activity against various fungi, such as *Fusarium oxysporum, Phytophthora infestans*, and *Botrytis cinerea*. Surfactin is believed to exert its antifungal effects by disrupting the fungal cell membrane and causing cell lysis [63]. Fengycin and iturin are thought to disrupt the fungal cell membrane and inhibit the biosynthesis of ergosterol, a component of fungal cell membranes, thereby exerting their antifungal effects [64].

The bacterial strains that showed significant PGP capacities in this study should be further tested *in vivo* for their potential of phytostimulation. The selected candidates will be m62a for its ability to solubilise phosphate, t2 for its high siderophore production, m72 for its ability to produce IAA and t5 for its ability to fix free nitrogen. These strains would be individually inoculated on model plants to characterise their effect on growth but also pooled together to test a formulation carrying all PGP capacities. *B. siamensis* related strains showed a strong ability to inhibit the growth of vanilla pathogens *in vitro*, as well as their ability to produce a wide range of lipoptides involved in inhibiting fungi and inducing systemic resistance in plants. Thus, the t24, m62b or m64 strains should be tested *in vivo*, not only on vanilla but also on other types of culture, as a natural phythoprotective agent.

## Ackowledgments

We would like to thank UMR PVBMT and Dr. Carine Charron for giving us the fungus strains of *C. orchidophilum* and *F. oxyspor*um used in this study.

## Author Contributions

HK designed and supervised the study. HK and FR sampled and isolated the bacteria strains. GLT, ZKC and HB contributed to the molecular biology and sequencing. AT, ZKC and HB performed experiments and analyzed the data for the PGP activities. GLT and JCM performed experiments for antagonisms assay. BB, PB and LQ performed experiments and analyzed the data for identification and distribution of lipopeptides. GLT prepared the original draft. BB and AT co-wrote the manuscript. HK, JCM and HB review and editing the manuscript. All authors have read and agreed to the published version of the manuscript.

## Supplementary Material

**Figure S1:**
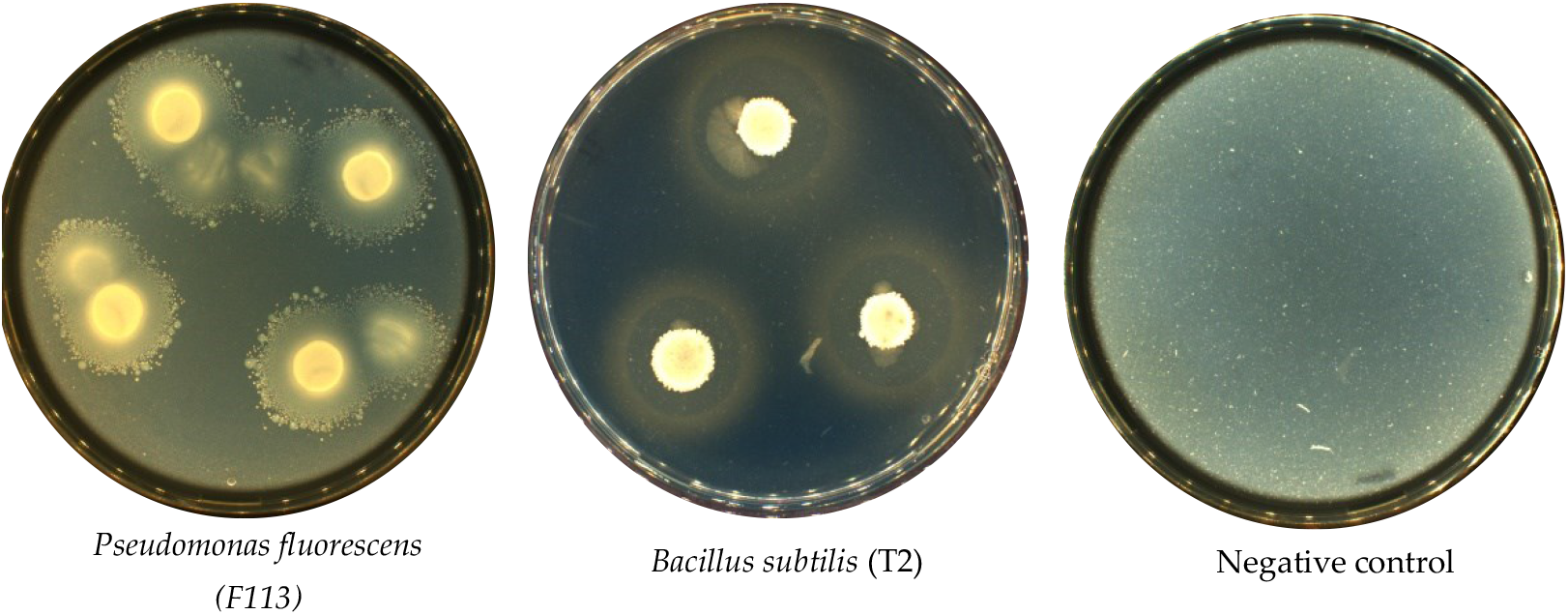
Halos produced by bacteria on MM9 – CAS agar by spot inoculation and incubation for 7 days at 28° C. (a) *Pseudomonas fluorescens* (F113) as positive control. (b) *Bacillus subtilis* (T2). (c) water as negative control. The halos around the colonies of bacteria indicating the ability of this isolate to extract siderophore that removes Fe from Fe-CAS agar medium.

**Figure S2:**
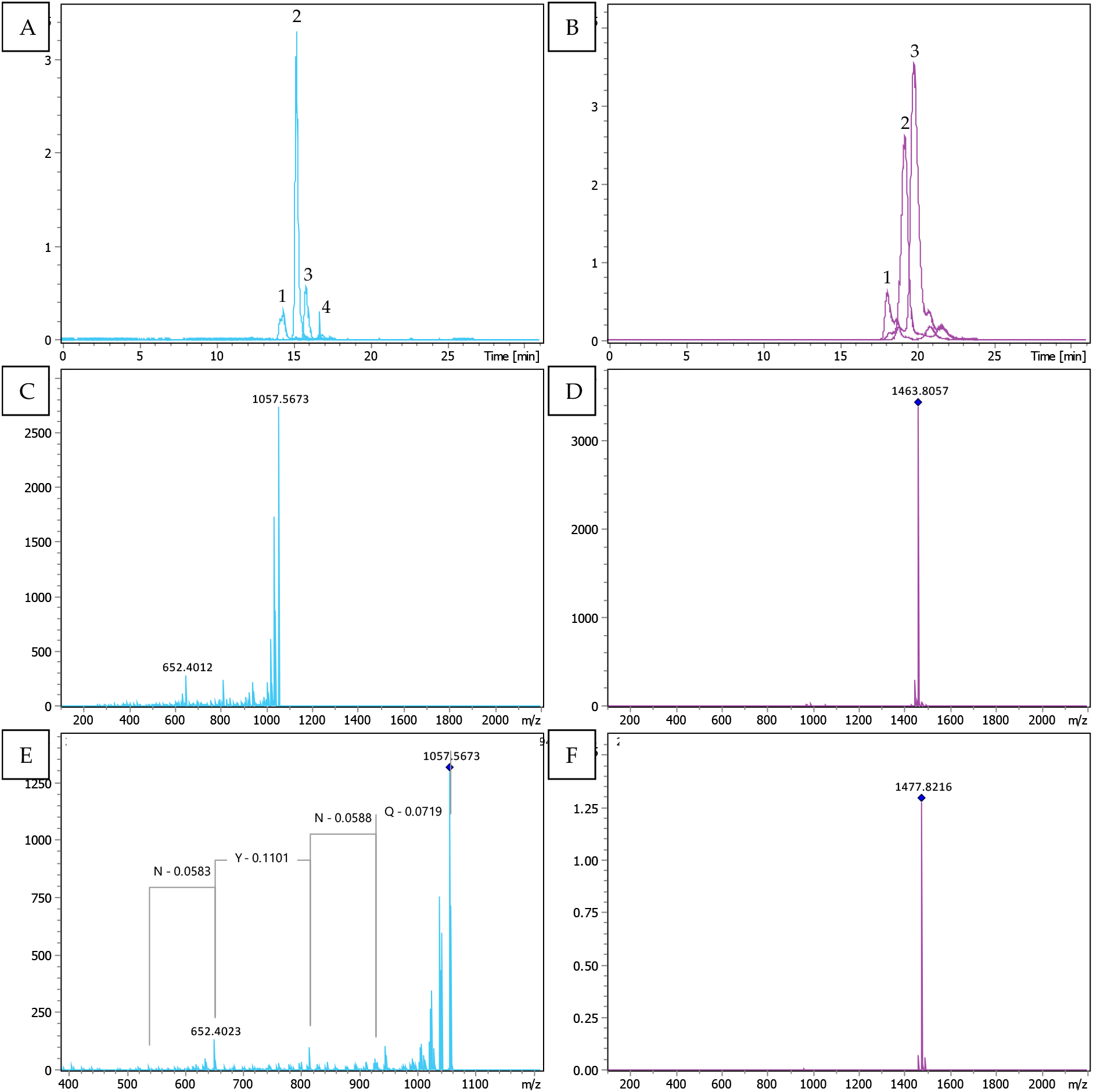
TIMS-TOF analysis of iturins and fengycins recorded in the inhibition area between *Bacillus siamensis* (m65) and *Fusarium oxysporum sp. Vanillae* (Fo72a). (A) The extracted ion chromatogram of 1043.5525 (1), 1057.56873 (2), 1071.5838 (3) and 1085.5995 (4) m/z representing the four present metabolites of iturins. (C) MS spectrum represented the single charged M+H+ ion of iturin 1057.56 m/z. (E) MSMS spectrum ion at 1057.56 m/z and the fragmentation lead to the peptide sequence analysis of iturin. (B) The extracted ion chromatogram of 1449.7891 (1), 1463.8057 (2) and 1477.8216 (3) m/z representing the three present metabolites of fengycin. (D) MS spectrum represented the single charged M+H+ ion of fengycin 1463.8057 m/z. (F) MS spectrum represented the single charged M+H+ ion of fengycin 1477.8216 m/z.

